# Quantification of reactive oxygen species production by the red fluorescent proteins KillerRed, SuperNova and mCherry

**DOI:** 10.1101/777417

**Authors:** John O. Onukwufor, Adam J. Trewin, Timothy M. Baran, Anmol Almast, Thomas H. Foster, Andrew P. Wojtovich

**Author notes:** Corresponding author: Andrew P. Wojtovich. These authors contributed equally (John O. Onukwufor, Adam J. Trewin).

## Abstract

Fluorescent proteins can generate reactive oxygen species (ROS) upon absorption of photons via type I and II photosensitization mechanisms. The red fluorescent proteins KillerRed and SuperNova are phototoxic proteins engineered to generate ROS and are used in a variety of biological applications. However, their relative quantum yields and rates of ROS production are unclear, which has limited the interpretation of their effects when used in biological systems. We cloned and purified KillerRed, SuperNova, and mCherry - a related red fluorescent protein not typically considered a photosensitizer - and measured the superoxide (O_2_ ^•-^) and singlet oxygen (^1^O_2_) quantum yields with irradiation at 561 nm. The formation of the O_2_ ^•-^-specific product 2-hydroxyethidium (2-OHE^+^) was quantified via HPLC separation with fluorescence detection. Relative to a reference photosensitizer, Rose Bengal, the O_2_ ^•-^ quantum yield (ΦO_2_ ^•-^) of SuperNova was determined to be 0.00150, KillerRed was 0.00097, and mCherry 0.00120. At an excitation fluence of 916.5 J/cm^2^ and matched absorption at 561 nm, SuperNova, KillerRed and mCherry made 3.81, 2.38 and 1.65 μM O_2_ ^•-^/min, respectively. Using the probe Singlet Oxygen Sensor Green (SOSG), we ascertained the ^1^O_2_ quantum yield (Φ^1^O_2_) for SuperNova to be 0.0220, KillerRed 0.0076, and mCherry 0.0057. These photosensitization characteristics of SuperNova, KillerRed and mCherry improve our understanding of fluorescent proteins and are pertinent for refining their use as tools to advance our knowledge of redox biology.

**GRAPHICAL ABSTRACT:** 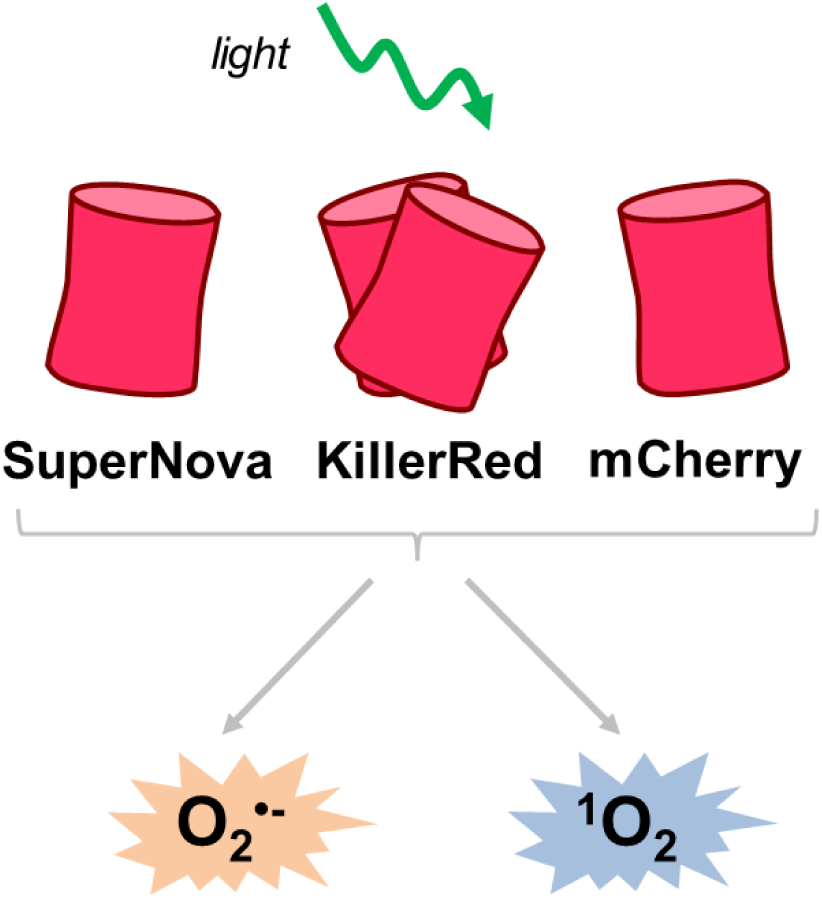

## INTRODUCTION

Fluorescent proteins generate reactive oxygen species (ROS) upon irradiation by type I or type II photosensitization mechanisms [1-4]. The type I mechanism involves electron transfer reactions that ultimately reduce molecular oxygen to form superoxide (O_2_ ^•-^) [3, 5]. Type II photosensitization involves the direct energy transfer from excited triplet state of the photosensitizer to oxygen to generate singlet oxygen (^1^O_2_) [4-7]. Both O_2_^•-^ and ^1^O_2_ can be formed by fluorescent proteins [4, 5] but the relative yields or fluxes depend on various factors, including the protein structure surrounding the chromophore, the oxygen concentration, temperature, and pH of the environment [3, 5].

A range of phototoxic fluorescent proteins have been developed such as KillerRed, KillerOrange, SuperNova, miniSOG and their derivatives; however their phototoxic properties are not fully characterized [1-3, 8-11]. KillerRed, a dimeric red fluorescent protein, was derived from a random and site-directed mutations of a jellyfish protein, anm2CP [1, 3, 10, 12]. KillerRed has a unique structure with a water channel to the chromophore that is responsible for its phototoxicity [1, 3, 10, 12]. The original KillerRed protein is prone to variable levels of dimerization, which can lead to artifacts and mislocalization of fusion proteins within a biological system [8]. These confounding factors can be mitigated by using the pseudo-monomeric version tandem KillerRed (tdKillerRed), which consists of two repeats of the KillerRed coding sequence, meaning that all copies are expressed as a dimer. SuperNova was derived from KillerRed and retains similar phototoxic properties but exists as a monomer, thereby limiting potential mislocalization events [8]. Both KillerRed and SuperNova are used in a variety of applications ranging from localized ROS production to cell ablation, however the quantities or the species of ROS responsible for the effect are often unclear. KillerRed has been used for chromophore-assisted light inactivation (CALI) in cells and organelles [1, 13-16]. These phototoxic effects have been shown to be sensitive to superoxide dismutase (SOD), catalase, and sodium azide [1, 8], suggesting that KillerRed possesses the capacity to generate both O_2_^•-^ (and subsequently hydrogen peroxide) and ^1^O_2_ oxidants [1, 2, 8]. Likewise, SuperNova has been shown to oxidize DHE and ADPA probes, implying that it too generates both O_2_ ^•-^ and ^1^O_2_ oxidants [8, 17].

Although the phototoxic effects of these fluorescent proteins to cellular functioning have been widely demonstrated, their precise ROS quantum yields, i.e. the ratio of ROS molecules generated per photon absorbed by the fluorophore, and intrinsic rates of ROS production have not previously been reported. Therefore, the aim of this study was to determine the quantum yields and rates of ROS production by phototoxic fluorescent proteins. Using Rose Bengal, a well-characterized chemical photosensitizer molecule with a defined O_2_ ^•-^ quantum yield (ΦO_2_ ^•-^) of 0.2 and ^1^O_2_ quantum yield (Φ^1^O_2_) of 0.75 as a standard [18], we determined the relative O_2_ ^•-^ and ^1^O_2_ quantum yields of KillerRed and SuperNova. As a negative control for photosensitization we used mCherry, a red fluorescent protein commonly used as an ‘inert’ fluorophore in many cellular imaging applications [3, 8]. Overall, we report the O_2_ ^•-^ and ^1^O_2_ quantum yield of the fluorescent proteins tdKillerRed and SuperNova, as well as mCherry.

## MATERIALS AND METHODS

### Protein cloning and purification

SuperNova, tdKillerRed, and mCherry were transformed and grown in a culture as previously described [8, 17]. SuperNova/pRSETB was a gift from Dr. Takeharu Nagai (Addgene plasmid # 53234) [8]. mCherry (pmCherry-C1) and tdKillerRed (#FP963, Evrogen) were amplified and ligated into pRSETB using BamHI and EcoRI. Plasmids were then transfected into JM109 (DE3) XJ autolysis cells, and protein expression was induced with isopropyl β-D-1-thiogalactopyranoside (IPTG). Cultures were centrifuged at 3200 g for 10 min, washed with PBS and flash frozen. Cell lysate was run through nickel beads, then protein was eluted with 100 µM imidazole in the presence of protease inhibitors (Roche) and desalted using a PD-10 column. Protein concentration was determined by Lowry assay, and absorbance scans were performed on a spectrophotometer (Shimadzu) to identify a region of spectral overlap in absorbance maxima between the proteins and Rose Bengal dye (# 330000, Sigma). The most robust overlap occurred between 550-580 nm (Fig. 1). Based on this, a 561 nm laser was chosen for subsequent experimentation. Proteins and Rose Bengal were diluted to achieve equal molar absorptivity at 561 nm using the Beer-Lambert equation (A=ε*b*c).

**Fig. 1.**
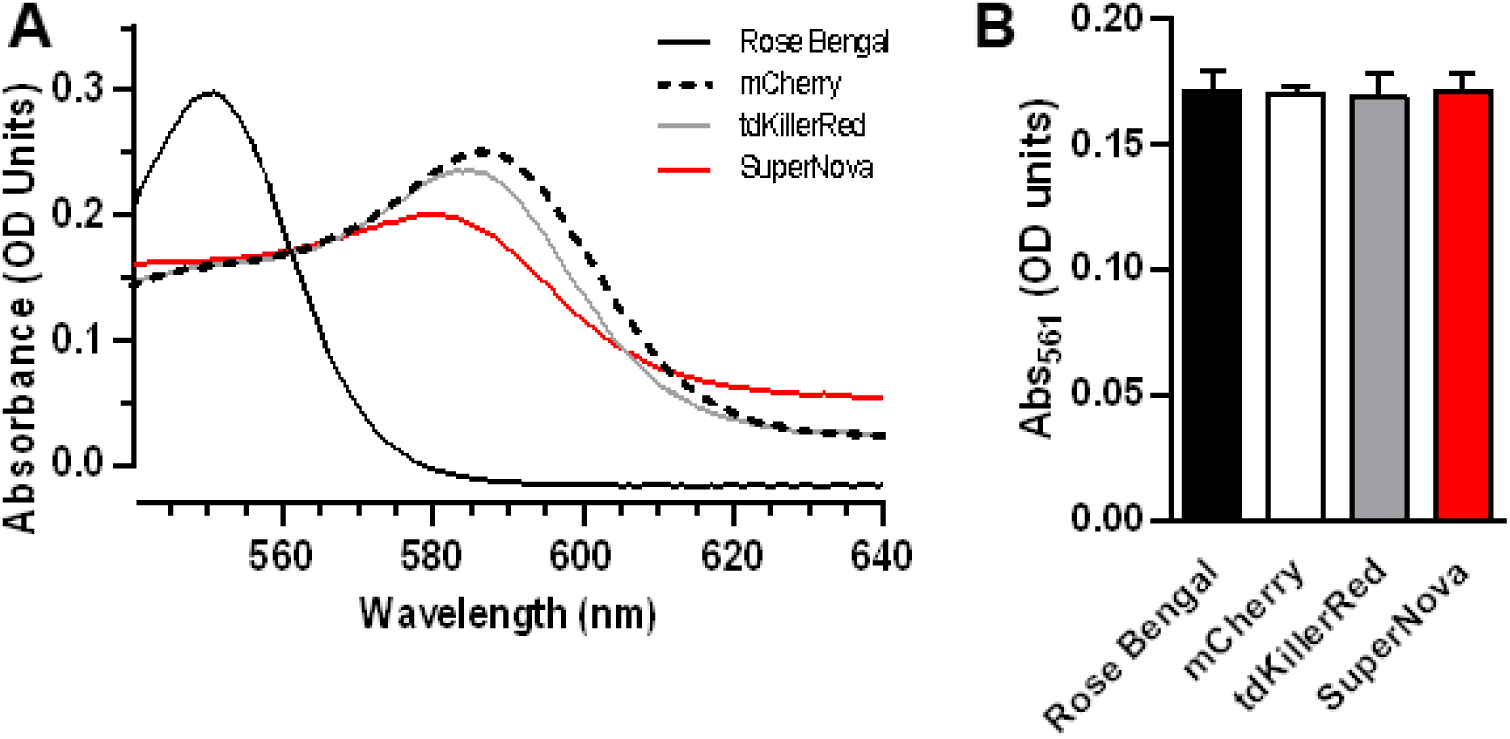
Equal photosensitizer absorbance at 561 nm. (a) Absorbance spectrum of photosensitizers. (b) Absorbance at 561 nm after adjustment of concentration of Rose Bengal dye (0.0026 mg/mL), mCherry (0.22 mg/mL), tdKillerRed (0.25 mg/mL), and SuperNova (0.76 mg/mL). Values are mean ± SD for n = 3 independent experiments; p > 0.05 by one-way ANOVA.

### Irradiation parameters

Irradiation of fluorescent proteins and the photosensitizing dye, Rose Bengal, were performed using a 561 nm class IIIb 50 mW diode laser (#1230935, Coherent® OBIS™, Edmund Optics, NJ, USA). The 0.7 mm diameter beam was focused through a 20x, 0.4 NA microscope objective lens (Swift) into a 200 µm core diameter, 0.22 NA SMA-terminated fiber optic cable (Part # M25L05, ThorLabs, Inc., Newton, NJ) for delivery to the sample. The fiber and objective lens were positioned using a Multimode Fiber Coupler Assembly (Part # F-91-C1-T, Newport Corporation, Irvine, CA). Fiber output was collimated with an aspheric lens (Part # A397TM-B, Thorlabs) to create a 2.5 mm-diameter collimated beam to irradiate each 200 µL sample volume contained within a 1.5 mL, 1 cm polystyrene cuvette (#97000-586, VWR). The irradiance was measured as 25 mW at the front surface of the sample cuvette using thermopile detector (818P-010-12, Newport Corporation, Irvine, CA) for all irradiation. Fluence/light dose (J/cm^2^) was modulated by adjusting irradiation time while maintaining a consistent fluence rate (mW/cm^2^).

### Determination of photobleaching rates

Photobleaching rates of photosensitizers (Rose Bengal, 0.0026 mg/ml; mCherry, 0.22 mg/ml; KillerRed, 0.25 mg/ml; SuperNova, 0.76 mg/ml) and the probe DHE alone and in combinations were determined in buffer (D-MRB; 220 mM Manitol, 70 mM Sucrose, 5 mM MOPS, 2 mM EGTA, 0.4% FFBSA, 0.1 mM DTPA, pH 7.3) at 20 °C. The fluorescence signal (Ex 525 nm; Em 550 nm) was acquired using a fluorescence spectrophotometer (Cary Eclipse, Agilent Technologies) during a cumulative time exposure (0-30 min) at 561 nm irradiation for determination of the reduction in fluorescence. To determine the bleaching rates with SOSG, DHE was replaced with SOSG in the buffer, and the change in absorbance was measured between 400 – 800 nm using a spectrophotometer.

### Xanthine oxidase superoxide production

Xanthine oxidase (XO) production of O_2_ ^•-^ was determined as the rate of SOD-sensitive cytochrome *c* reduction, as previously described [7, 19]. Briefly, XO (0.25, 0.50, 1.0 and 4.0 mU/mL) was added to a 1 cm cuvette containing cytochrome *c* (40 µM) in PBS containing DTPA (D-PBS: 7.78 mM Na_2_HPO_4_, 2.20 mM KH_2_PO_4_, 0.1 mM DTPA, pH 7.3). All reactions were carried out at ambient O_2_ and where indicated catalase (4200 U/mL) or SOD (800 U/mL) was present. Baseline was collected for 2 min before 1 mM of xanthine (X) was added to initiate the reaction. Cytochrome *c* reduction was monitored at 550 nm for 10 min, and the rate was calculated using an extinction coefficient of 18.7 mM^-1^ cm^-1^ [20^].^

### Superoxide quantification

The oxidation of dihydroethidium (DHE) yields the O_2_ ^•-^ specific fluorescence product 2-hydroxyethidium (2-OHE^+^) along with non-specific fluorescent products including ethidium (E^+^), which were separated using HPLC as previously described [7, 17, 21, 22]. Briefly, XO (4 mU/mL) and X (1 mM) were incubated in D-PBS at 20 °C for the indicated time (0 – 60 min). Rose Bengal (0.0026 mg/mL), mCherry (0.22 mg/ml), tdKillerRed (0.25 mg/ml), or SuperNova (0.76 mg/ml) were irradiated at 561 nm for the indicated time (0 – 30 min) in D-MRB in the presence of DHE (100 μM). For experiments containing photosensitizers, the absorbance was measured (400-800 nm) both pre- and post-irradiation at 561 nm. To these samples, an equal volume of 200 mM HClO_4_/MeOH was added, centrifuged at 17,000 x *g*, and the supernatant transferred to an equal volume 1 M K^+^PO^4 -^ at pH 2.6.

Samples were separated using a Polar-RP column (Phenomenex, 150 x 2 mm; 4µm) on a Shimadzu HPLC with fluorescence detection (RF-20A). The flow rate was constant (0.1 mL/min) using a gradient of two mobile phases (A: 10% ACN, 0.1 %TFA; B: 60% ACN, 0.1 %TFA). The gradient was the following: 0 min, 40% B; 5 min, 40% B; 25 min, 100% B; 30 min, 100% B; 35 min, 40% B; 40min, 40% B. Standard curves were generated against known concentrations of E^+^ and 2-OHE^+^, and peaks were quantified using Lab Solutions (Shimadzu) [7, 17].

### Singlet oxygen quantification

The ^1^O_2_ production of photosensitizers (Rose Bengal, 0.0026 mg/mL; mCherry, 0.22 mg/ml; KillerRed, 0.25 mg/ml; SuperNova, 0.76 mg/ml) was measured using SOSG (1 µM, #S36002, Invitrogen) in D-MRB at 20°C [7]. The SOSG signal (Ex 525 nm; Em 550 nm) was acquired using a Cary Eclipse fluorescence spectrophotometer (Agilent Technologies) pre- and post-561 nm irradiation for determination of the change in SOSG fluorescence intensity [7].

### Calculations and statistical analysis

Fluorescent protein O_2_ ^•-^ and ^1^O_2_ quantum yields were determined after correcting for the bleaching rates of the photosensitizers, as we have previously demonstrated the importance of photobleaching in explaining time-dependent ROS production by photosensitizers [23]. Measurements of fluorescence/absorbance vs. illumination duration were first normalized to the value prior to illumination, and then fit with an equation of the form *B*(*t*) = *ae*^−*bt*^, where *a* and *b* are fit coefficients and *t* is the illumination duration in seconds. The total number of absorbed photons for a sample can then be expressed as 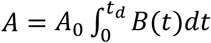, where A_0_ is the absorption prior to illumination, *t*_*d*_ is the illumination duration, and *B(t)* is the bleaching curve described above. Relative to a reference quantum yield (Φ_R_), the quantum yield of a sample (Φ_S_) can be determined by 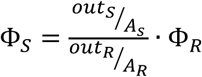, where *out* is the output of interest and *A* is the total number of absorbed photons, as described above. Incorporating correction for bleaching of the sample and reference, with knowledge that pre-illumination (*A*_*0*_) is equal for all samples, the quantum yield can be expressed as:

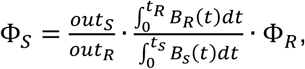

where *out*_*S*_ *and out*_*R*_ are measured outputs for illumination durations of *t*_*S*_ and *t*_*R*_ for the sample and reference, respectively, and *B*_*s*_ and *B*_*R*_ are the corresponding bleaching curves. All fitting and calculation was performed in MATLAB (The MathWorks, Inc., Natick, MA).

O_2_ ^•-^ production rates of the fluorescent proteins were calculated based upon the standard curve generated using X/XO. Since the apparent number of O_2_ ^•-^ molecules required to generate one 2-OHE^+^ molecule is dependent on the rate of O_2_ ^•-^, photosensitizer O_2_ ^•-^ production was matched with 4 mU/mL XO superoxide generation. Under these conditions X/XO produced 2.24 µM O_2_ ^•-^/min. X/XO was incubated (0-60 min) of DHE and 2-OHE^+^ was measured and plotted against the expected cumulative O_2_ ^•-^ concentration generated during that time, as previously described [7]. At these lower rates the ratio of O_2_^•-^ to 2-OHE^+^ was linear (y = 55.62(x) + 326.2; R^2^ = 0.98).

Statistical analysis: Data were first tested for normality of variance, and were then analyzed by one- or two-way ANOVA with Tukey’s post hoc using GraphPad Prism (v7).

## RESULTS

### Purification and characterization of fluorescent proteins

Fluorescent proteins subjected to SDS-PAGE migrated at their expected molecular weight (Supplemental Fig. 1). In order to measure protein photosensitization characteristics relative to a reference dye (Rose Bengal), we first sought to determine i) a wavelength that was near the absorption maxima for each chromophore, ii) a concentration of each chromophore in solution that would allow all of the photosensitizers absorb an equal number of photons and iii) is not confounded by absorption of photons by other reagents used for detection of ROS. We determined from absorbance spectra that excitation at 561 nm met each of these criteria (Fig. 1A), and photosensitizer concentrations were then optically matched for equal absorbance at 561 nm (Fig. 1B).

### Superoxide quantum yield and superoxide generation rate of fluorescent proteins

We measured light-dependent photosensitizer O_2_ ^•-^ generation using HPLC to quantify 2-OHE^+^, a O_2_ ^•**-**^ specific reaction product of DHE [7, 24-26]. Since the known yield of Rose Bengal served as our reference, we confirmed that Rose Bengal produced 2-OHE^+^ in a light dose-dependent manner (Fig. 2A). Similarly, the fluorescent proteins tdKillerRed, SuperNova and mCherry also produced 2-OHE^+^ in a light dose-dependent manner (Fig. 2B), yet the magnitude of 2-OHE^+^ for the protein photosensitizers was considerably lower than that of Rose Bengal. For example, after 60 seconds of illumination Rose Bengal generated ∼17,000 pmol/mL 2-OHE^+^, while after 300 seconds the fluorescent proteins produced ∼500 pmol/mL (Fig. 2B).

**Fig. 2.**
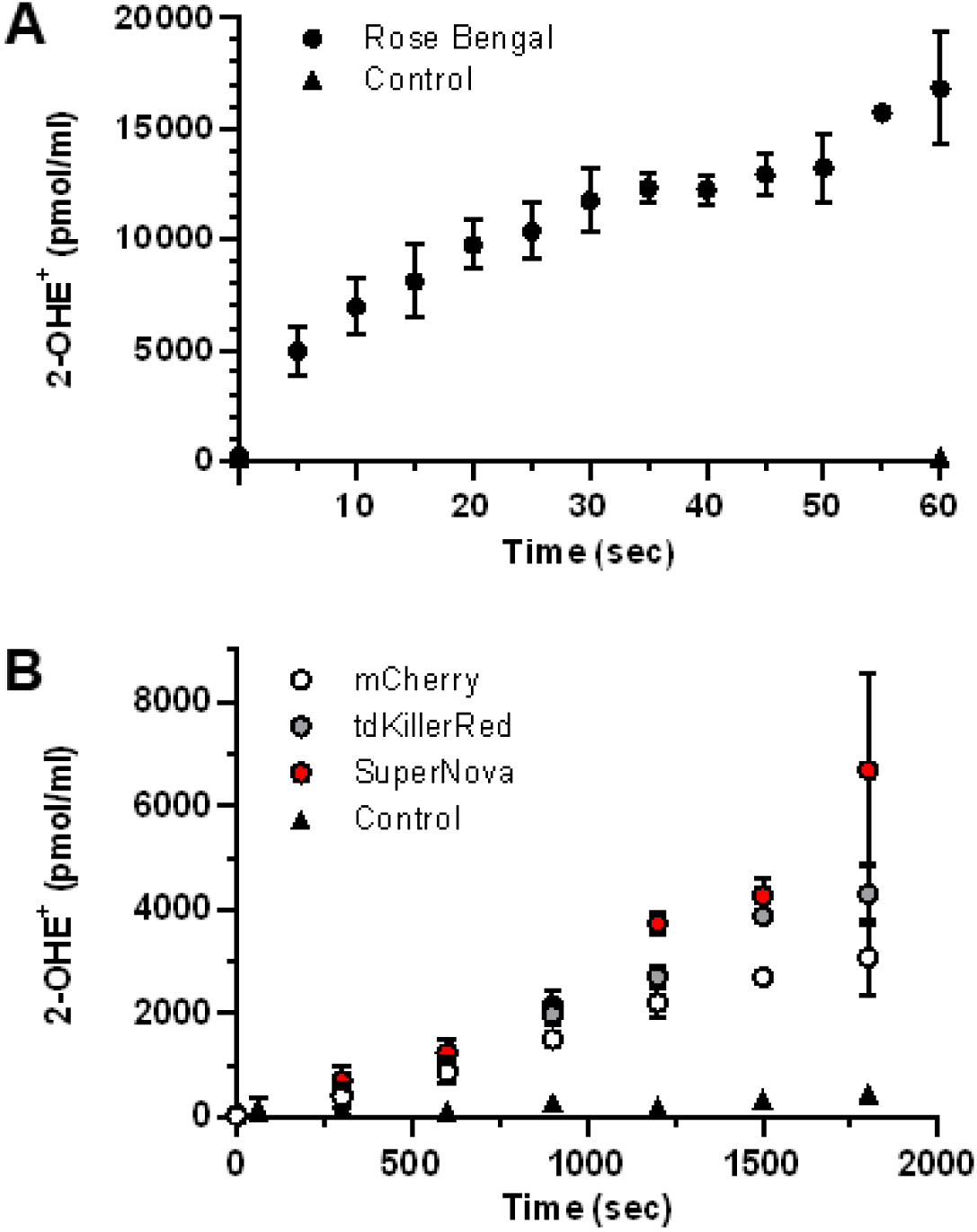
Light-dependent superoxide generation by photosensitizers. (a) Rose Bengal, (b) tdKillerRed, Supernova, mCherry and control (no photosensitizer) were irradiated with equal molar absorptivity at 561 nm in the presence of DHE (100 µM) for quantification of 2-OHE^+^. Values are mean ± SD for n = 3 independent experiments.

Next, we sought to determine the O_2_ ^•**-**^ quantum yield of tdKillerRed, SuperNova and mCherry relative to Rose Bengal, with a known ΦO_2_ ^•-^ of 0.2 [18]. The determination of quantum yields relies on the equal absorbance of photons, yet photobleaching results in a decrease in photon absorbance over time that occurs at different rates between photosensitizers. We therefore measured the rate of photosensitizer bleaching by assessing the change in fluorescence in response to a cumulative light-dose. We then corrected for the bleaching rates of the individual fluorophores and the probe, DHE, (Supplemental Fig. S2) in order to calculate the ΦO_2_ ^•-^ relative to Rose Bengal. We thus determined that SuperNova had a ΦO_2_ ^•-^ of 0.0015, and tdKillerRed’s ΦO_2_ ^•-^ was 0.00097; mCherry had a comparable ΦO_2_ ^•-^ (Table 1).

**TABLE 1:** Superoxide and singlet oxygen quantum yield of mCherry, tdKillerRed, and SuperNova.

| Fluorescent proteins | $O_2^{\cdot-}$ Quantum yield ($\phi O_2^{\cdot-}$) | $^1O_2$ Quantum yield ($\phi^1O_2$) |
| --- | --- | --- |
| mCherry | 0.00120±0.000044 | 0.0057±0.00027 |
| tdKillerRed | 0.00097±0.000042 | 0.0076±0.00026 |
| SuperNova | 0.00150±0.000016 | 0.0220±0.00180 |

Data are mean ± SD for n=3 independent experiments

We next sought to calculate the O_2_ ^•-^ production rate of fluorophores. However, the apparent ratio of O_2_ ^•-^ molecules necessary to form one molecule of 2-OHE^+^ is highly dependent of the rate of O_2_ ^•-^ generation, possibly due to competition with spontaneous dismutation [7, 26]. Therefore, we generated a standard curve using a concentration of xanthine oxidase that produces O_2_ ^•-^ at a similar rate to that of the photosensitizers. Based on the results of the dose response (Supplementary Fig. S3), we selected 4 mU/mL of xanthine oxidase (Fig. 3A) to match the 2-OHE^+^ production rates from our photosensitizers at this concentration and light dose. We determined that 4 mU/mL of xanthine oxidase produces 2.44 µM/min of O_2_ ^•-^, which was SOD-sensitive and catalase-insensitive (Fig. 3A). We incubated the same amount of xanthine oxidase in the presence of DHE and measured the formation 2-OHE^+^ over time and expressed it as a function of expected cumulative O_2_^•-^ production. Our results show a linear increase of 2-OHE^+^ with increasing amounts of O_2_ ^•-^ across the tested range (R^2^ = 0.98; Fig. 3B). Given that the photosensitizers absorbed an equal amount of light and hence have the same ability to make ROS (Fig. 1), we then used this equation to derive the rate of O_2_ ^•-^ production by photosensitizers per unit light dose (Fig. 3C & D) from the data in Fig. 2. Rose Bengal had the highest rates of O_2_ ^•-^ production across light doses (∼300 µM O_2_ ^•-^ /min at 30.55 J/cm^2^) with mCherry producing the least amount O_2_ ^•-^ per light dose (∼1.65 µM O_2_ ^•-^ /min at 916.5 J/cm^2^) (Fig. 3C & D). The rate of O_2_ ^•-^ production by Rose Bengal decreased with increasing light dose, which is consistent with the bleaching rate of Rose Bengal (Supplemental Fig. 2). The progressive loss of absorption resulted in a fluence-dependent decrease in the O_2_ ^•-^ production rate. At the light doses tested, each of the fluorescent proteins showed an increasing O_2_ ^•-^ production rate that reached a plateau around 600 J/cm^2^ (Fig. 3D). The gradual increase in the measured O_2_ ^•-^ production rate could be due to saturation of local reaction sites, such as amino acids, that can quench ROS, or a conformational change in the protein resulting in a maximal observed production rate, as has been reported for other fluorescent proteins [27]. As the fluorescent proteins bleach at a slower rate compared to Rose Bengal (Supplemental Fig. 2), we did not observe the same decrease in O_2_^•-^ production rate that was observed for Rose Bengal.

**Fig. 3.**
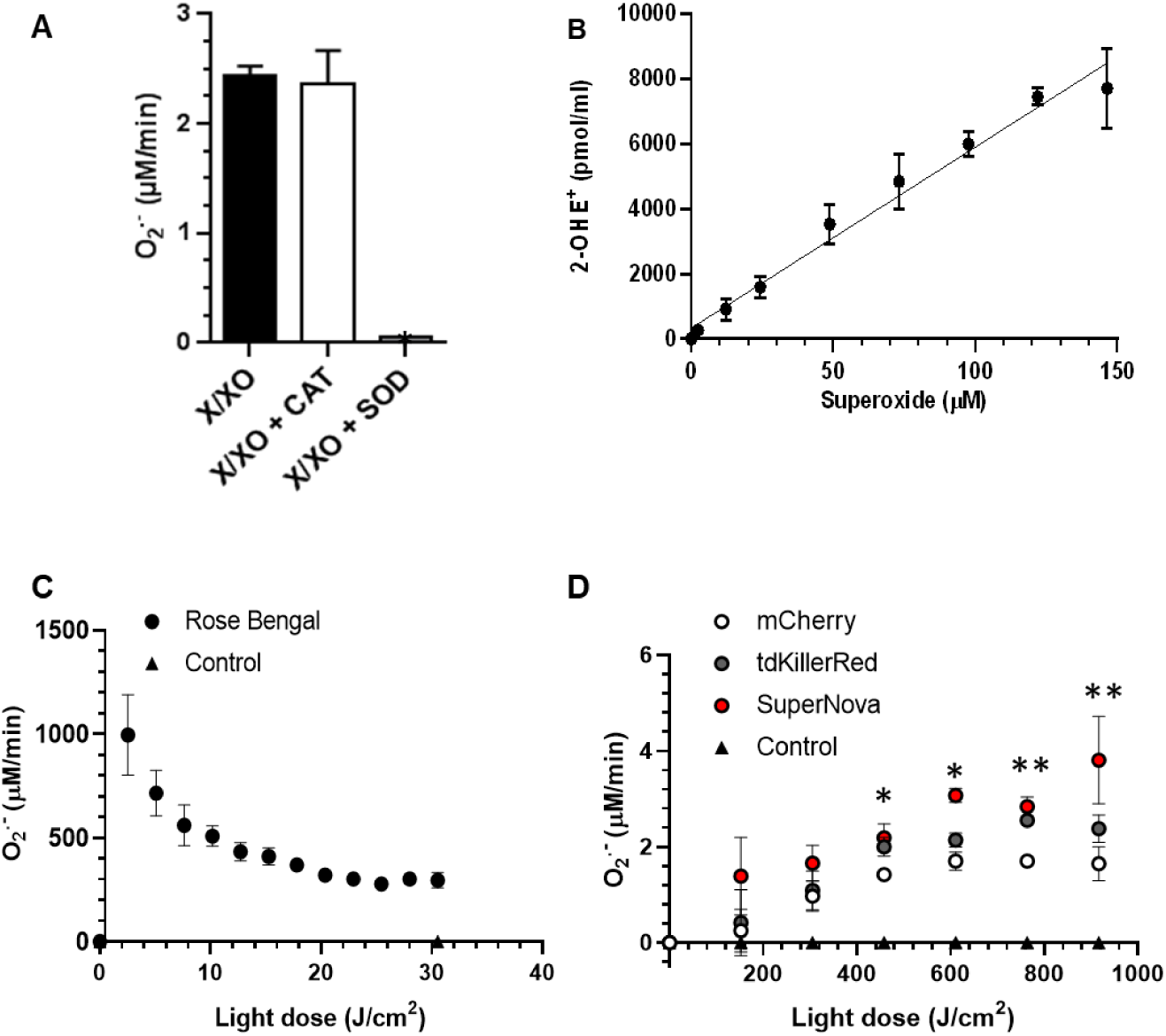
Determination of superoxide production per light dose. (a) xanthine/xanthine oxidase (X/XO) O_2_^•-^ production was assessed using cytochrome c reduction assay. Xanthine oxidase (XO, 4 mU/mL) and xanthine (X, 1 mM) were incubated with catalase (CAT) superoxide dismutase (SOD) where indicated. *p < 0.05 vs X/XO and X/XO+CAT, one-way ANOVA, Tukey post hoc. (b) Time course (0-60 min) of X/XO O_2_^•-^ generation was measured using HPLC separation of 2-OHE^+^ and then plotted against the expected O_2_^•-^ production. (c) Rose Bengal O_2_^•-^production rate per light dose. (d) Fluorescent protein (mCherry, tdKillerRed and SuperNova) O_2_^•-^ production per light dose. Data from (c) and (d) are derived from data presented in Fig. 2. *p < 0.05 SuperNova vs mCherry, ** p < 0.05 SuperNova and tdKillerRed vs mCherry, two-way ANOVA, Bonferroni post hoc. Values are mean ± SD for n = 3 independent experiments.

### Singlet oxygen quantum yield of fluorescent proteins

Singlet oxygen sensor green (SOSG) specifically detects ^1^O_2_ [4, 7, 27, 28] and does not react with other ROS, such as O_2_ ^•-^ or the hydroxyl radical, making it a suitable ^1^O_2_ detector under conditions were multiple ROS are being generated [29]. We assessed the ^1^O_2_ production of the photosensitizers by measuring the relative change of SOSG fluorescence and correcting for the bleaching rate of the individual fluorophores (Supplemental Fig. 2) [18]. Rose Bengal had the greatest SOSG fluorescence change with irradiation time (Fig. 4A) relative to those of the fluorescent proteins (Fig. 4B). The Φ^1^O_2_ of the fluorescent proteins were then calculated relative to the Rose Bengal reference Φ^1^O_2_ of 0.75 [18]. We determined that SuperNova had the highest Φ^1^O_2_ at ∼0.022, while mCherry had the lowest Φ^1^O_2_ of ∼0.0057 (Table 1). This demonstrates that mCherry, KillerRed and SuperNova are each capable of generating ^1^O_2_ in an irradiation dose dependent manner.

**Fig. 4.**
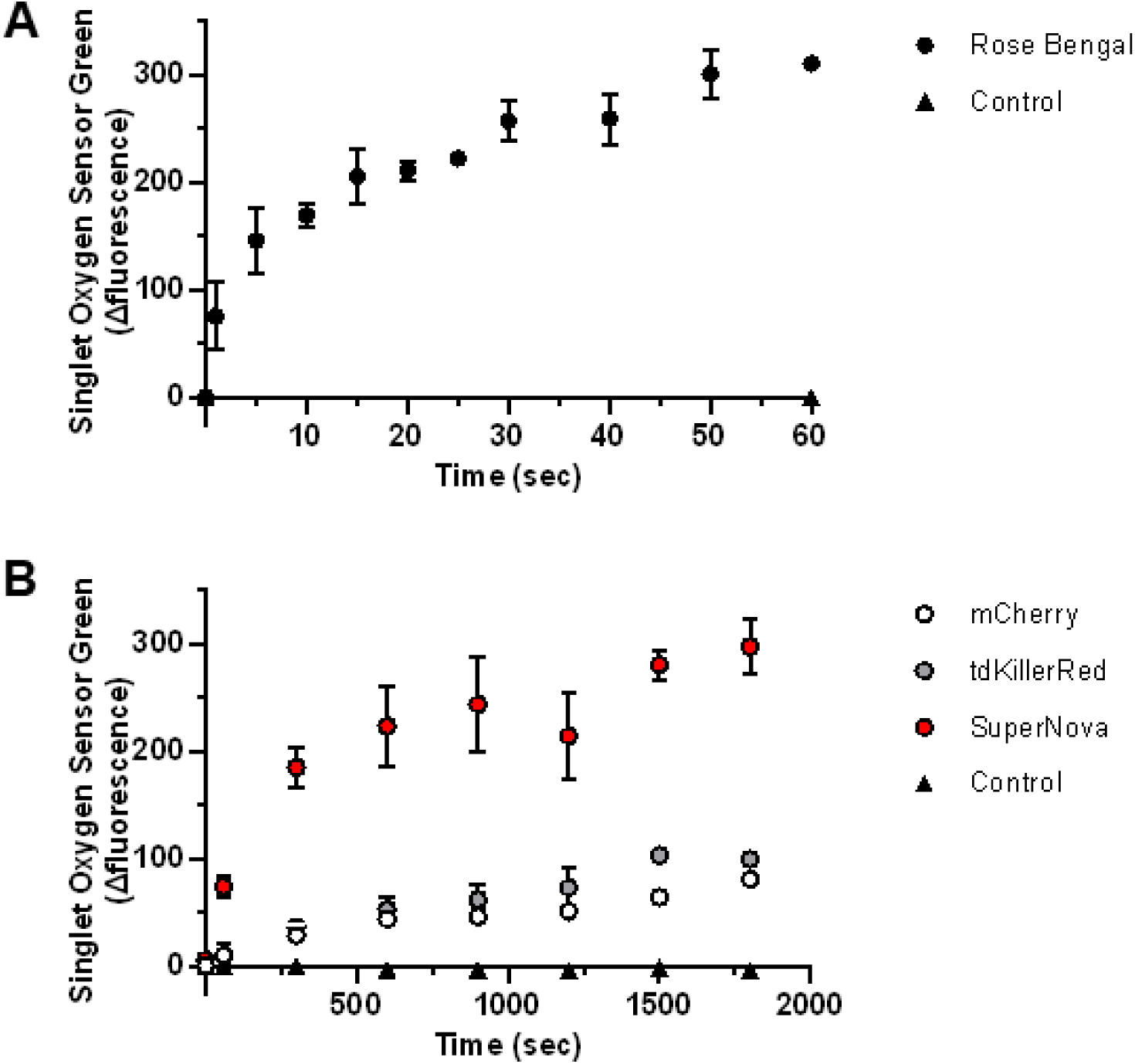
Singlet oxygen generation by photosensitizers in response to 561 nm irraditation. (a) Rose Bengal, (b) tdKillerRed, Supernova, mCherry and control (no photosensitizer) were irradiated with equal molar absorptivity at 561 nm in the presence of 0.1 µM SOSG. The initial fluorescence reading (Ex 525 nm; Em 550 nm) was subtracted from the post-illumination reading and presented as the relative fluorescence change. Values are mean ± SD for n = 3 independent experiments.

## DISCUSSION

The main findings from this study are that the red fluorescent proteins tdKillerRed, SuperNova, and mCherry each generate O_2_ ^•-^ and ^1^O_2_ via type I and II mechanisms, respectively. We also report for the first time quantitative ROS quantum yields for tdKillerRed, SuperNova and mCherry fluorescent proteins.

Genetically-encoded photosensitizers are used in a variety of biological applications to generate ROS in a light-dependent manner. They have the advantage of being targeted to precise regions in the cell to provide spatial control over ROS production [2]. However, their precise ROS producing characteristics are often overlooked provided that a biological phenotype has been observed. In contrast to Φ^1^O_2_, very little is known about fluorescent protein ΦO_2_ ^•-^. This may be a result of the limited methods to selectively detect O_2_ ^•-^, although one study has reported the ΦO_2_ ^•-^ and Φ^1^O_2_ of red fluorescent protein, TagRFP [4]. Using a similar SOSG detection approach, the Φ^1^O_2_ was estimated at 0.004, while the ΦO_2_ ^•-^ was estimated at <0.0002 using DHE bleaching as a measure of O_2_ ^•-^ [4].

The first developed photosensitizer protein, KillerRed, was initially reported to make O_2_ ^•-^ and ^1^O_2_ [1, 10]. Subsequently, literature has suggested that the KillerRed photosensitization mechanism selectively produces O_2_ ^•-^ and relies on the water channel to the chromophore for its phototoxicity [10, 30]. Depending on the application, one type of ROS may predominate in contributing to the light-induced effect. For example, ^1^O_2_ played a role in KillerRed CALI experiments [1], while O_2_ ^•-^ mediated phototoxicity [31]. While our results demonstrate that both ^1^O_2_ and O_2_ ^•-^ are capable of being produced, researchers should consider which species is relevant to their particular biological application.

SuperNova was derived from KillerRed, and it would be reasonable to assume that the photosensitization mechanisms would be similar. Accordingly, SuperNova has been thought to produce O_2_ ^•-^ and ^1^O_2_, as measured by 2-OHE^+^ formation [17] and ADPA photobleaching [8], respectively. In the present study, SuperNova’s comparatively larger ROS quantum yield than KillerRed is consistent with previous reports of greater phototoxicity [8]. Specifically, at 916.5 J/cm^2^ of fluence, we show that the SuperNova O_2_ ^•-^ production rate is ∼1.6 fold higher than KillerRed. However, the O_2_ ^•-^ production rate was not consistent across fluences tested, and plateaued at the highest light dose tested. While the quantum yields provide a direct comparison of the phototoxic mechanisms of the red fluorescent proteins tested, caution is warranted when extrapolating these findings *in vivo*. The O_2_ and pH gradients or endogenous chromophores present in the cellular milieu can affect the ROS generation by photosensitizers. For example, a high O_2_ tension could favor ^1^O_2_ production, while hypoxic conditions could favor O_2_ ^•-^ production [32].

Unlike KillerRed and SuperNova that were derived from the jellyfish protein anm2CP, mCherry was derived from the sea anemone protein DsRed. Owing to the structural differences that exist due to their independent lineage, mCherry lacks a water channel, suggesting that it would not be as phototoxic as KillerRed. Indeed, it is widely used in biological applications under the assumption that it is photochemically inert. However, some previous reports have also shown that mCherry can be phototoxic [33] and that it produces O_2_ ^•-^ [8^, 34]^ and ^1^O_2_ [8] upon irradiation. Our present findings are in agreement with this and indicate that mCherry actually displays similar ΦO_2_ ^•-^ and Φ^1^O_2_ as ‘professional’ photosensitizer proteins.

The genetically-encoded photosensitizers display ΦO_2_ ^•-^ and Φ^1^O_2_ that are orders of magnitude lower than the chemical photosensitizer Rose Bengal. Yet, despite their lower quantum yields, their ability to generate a biologically relevant effect is well established [1, 2, 8, 17]. Once formed by the excited chromophore, the superoxide anion must escape the protein barrel structure in order to be released to the surrounding environment and react with the ROS probe [10]. The protein barrel likely shields the release of ROS, potentially explaining the lower observed ΦO_2_ ^•-^ of the protein photosensitizers compared to Rose Bengal which can directly release oxidants to the surrounding aqueous environment. Nevertheless, our current Φ^1^O_2_ findings are generally in agreement with other fluorescent proteins that have been reported to range from Φ^1^O_2_ 0.004 to 0.030 [3]. Recently, optimized variants have reportedly reached Φ^1^O_2_ ∼0.6 [35]. New approaches are aimed at combining the large quantum yields of chemical photosensitizers with the advantages of genetically-encoded photosensitizers [36].

### Conclusion

Overall, we demonstrate that the red fluorescent proteins tdKillerRed, SuperNova and mCherry are able to photosensitize O_2_ ^•-^ and ^1^O_2_. Our studies provide ΦO_2_ ^•-^, Φ^1^O_2_ and rates of O_2_ ^•-^ production across light doses. Our findings will help elucidate mechanisms mediated by phototoxic proteins and aid in the development of efficient or selective ROS production by genetically-encoded photosensitizers [35, 37].

## ABBREVIATIONS

2-OHE^+^: 2-hydroxyethidium
CALI: Chromophore assisted light inactivation
DHE^+^: Dihydroethidium
E^+^: Ethidium
O_2_ ^•-^: Superoxide
^1^O_2_: Singlet oxygen
Φ: Quantum yield
ΦO_2_ ^•-^: Superoxide quantum yield
Φ^1^O_2_: Singlet oxygen quantum yield
ROS: Reactive oxygen species
SOSG: Singlet oxygen sensor green
SOD: Superoxide dismutase
X: Xanthine
XO: Xanthine oxidase

## ACKNOWLEDGEMENTS

We thank the members of the mitochondrial research group at the University of Rochester Medical Center for their valuable suggestions and contributions. AJT current address: Institute for Physical Activity and Nutrition (IPAN), School of Exercise and Nutrition Sciences, Deakin University, Burwood, VIC, Australia.

---

**Supplementary Fig. 1.**
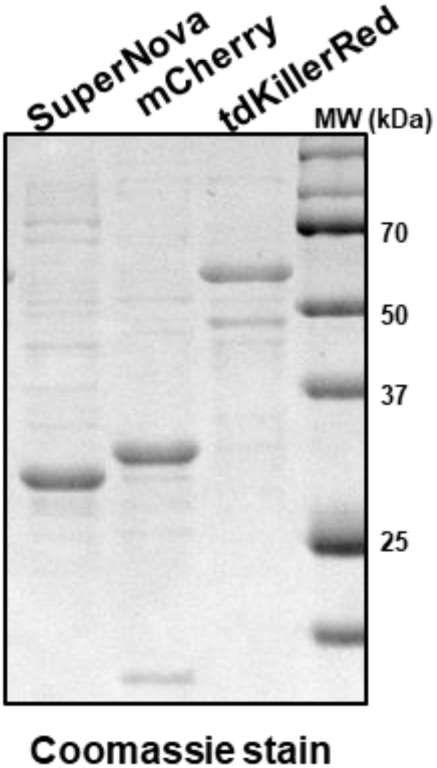
Purified fluorescent proteins. Purified SuperNova, mCherry, and tandem KillerRed protein were denatured in buffer containing 100 mM Tris HCl, 4% w/v SDS, 10% v/v glycerol, 0.2% w/v bromophenol blue, 2% v/v β-mercaptoethanol then heated for 5 min at 95°C before separation by SDS-PAGE (10 µg per lane) and then stained with coomassie blue.

**Supplementary Fig. 2.**
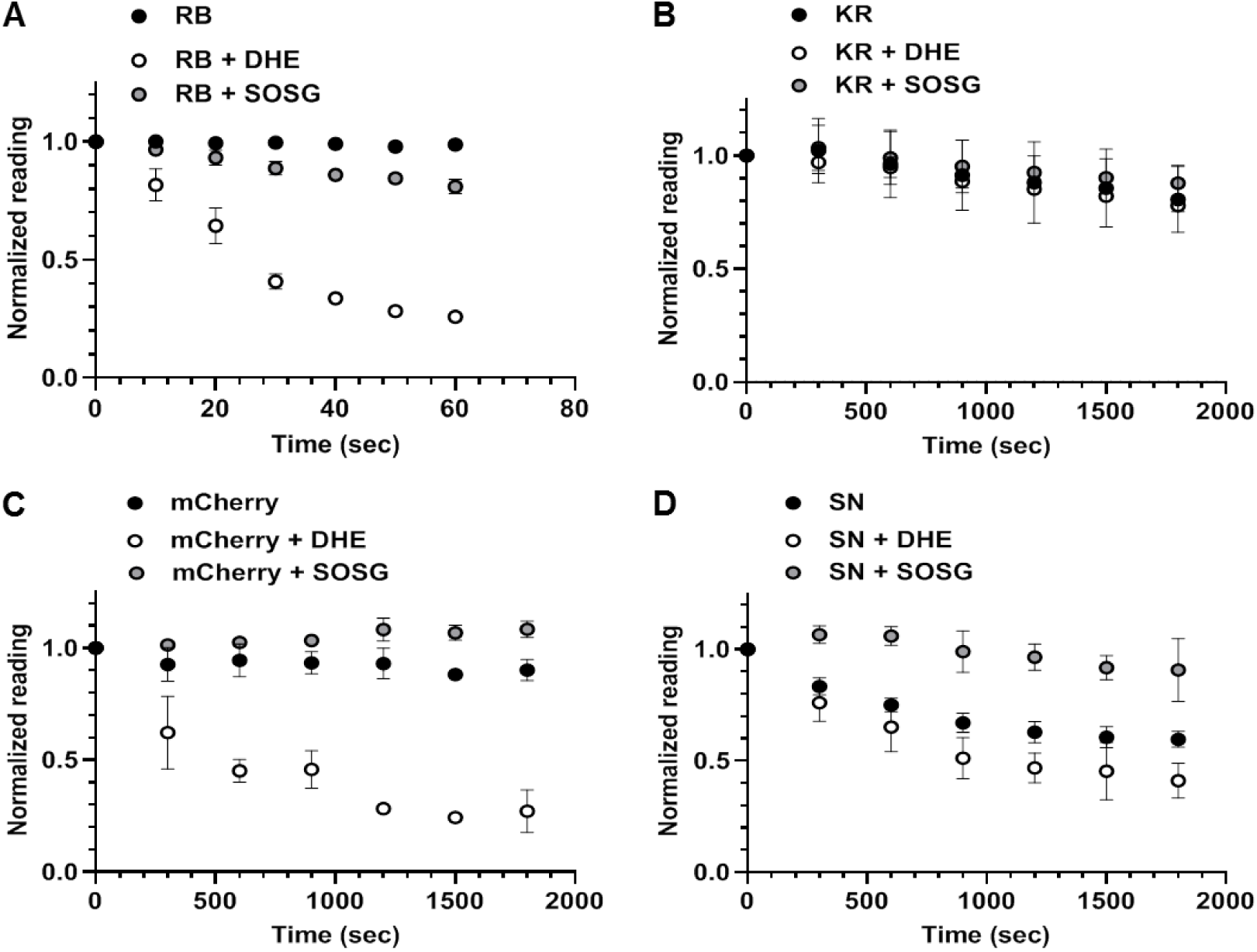
Photobleaching rates of photosensitizers with DHE or SOSG. (A) Rose Bengal, (B) KillerRed, (C) mCherry, and (D) SuperNova alone or in the presence of dihydroethidium (DHE) or Singlet Oxygen Sensor Green (SOSG). The cumulative fluorescence (photosensitizer alone or with DHE) or cumulative absorbance (photosensitizer with SOSG) was measured (0-60 sec for Rose Bengal; 0-30 min for KillerRed, mCherry, and SuperNova) following irradiation at 561 nm. Data were then normalized to baseline after bleaching fit correction using MATLAB. Data are N = 3, mean ± SD.

**Supplementary Fig. 3.**
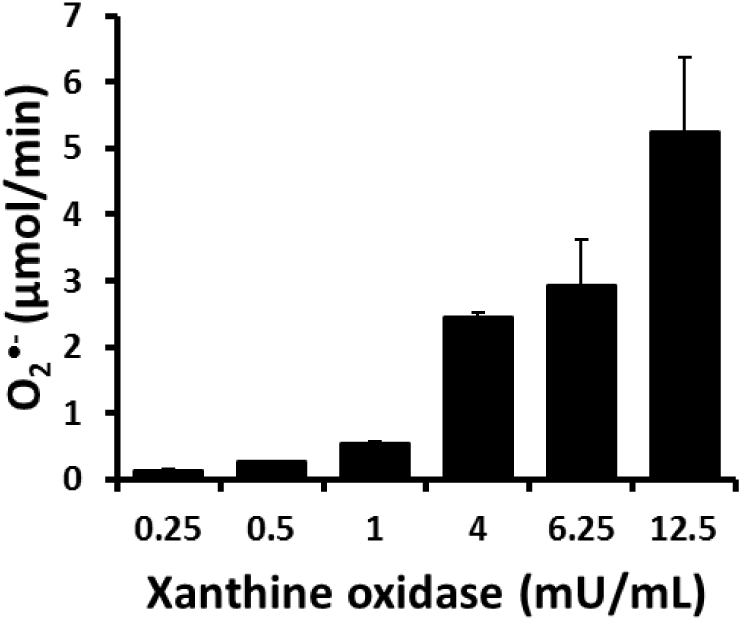
Dose dependent superoxide production of xanthine oxidase. Xanthine oxidase (0.25, 0.5, 1, 4, 6.25 and 12.5 mU/mL) superoxide production was assessed with in the presence xanthine (1mM) using cytochrome c reduction. Data are N = 4 independent values, mean ±SD.

